# Morphology-dependent entry kinetics and spread of influenza A virus

**DOI:** 10.1101/2024.08.01.605992

**Authors:** Sarah Peterl, Carmen M. Lahr, Carl N. Schneider, Janis Meyer, Xenia Podlipensky, Vera Lechner, Maria Villiou, Larissa Eis, Steffen Klein, Charlotta Funaya, Elisabetta Ada Cavalcanti-Adam, Frederik Graw, Christine Selhuber-Unkel, Karl Rohr, Petr Chlanda

## Abstract

Influenza A viruses (IAV) display a broad variety of morphologies ranging from spherical to long filamentous virus particles. These diverse phenotypes are believed to allow the virus to overcome various immunological and pulmonary barriers during entry into the airway epithelium and influence the viral entry pathway. Remarkably, lab-adapted IAV strains lost this morphological variance and exhibit preferred spherical morphology. However, it remains unclear which factors lead to this lab-adapted preference and which pulmonary defense factors are responsible for the preferred filamentous morphology in physiological settings. In this study, we established fluorescent reporter viruses with spherical or filamentous morphology but with the same surface glycoproteins. We developed a correlative fluorescence and scanning electron microscopy workflow to analyze the impact of viral morphology on cell-to-cell spread and identify conditions under which IAV with either spherical or filamentous morphology confer an advantage. Our findings demonstrate that filamentous IAV cell-to-cell spread is significantly slower in various cell lines, which can explain the predominant spherical morphology in lab-adapted strains. This observation is consistent with delayed entry kinetics of filamentous viruses structurally analyzed by cellular cryo-electron tomography. We found that cellular junction integrity and mucin do not exert morphology-dependent inhibition of IAV cell-to-cell spread. On the other hand, filamentous virions confer an advantage under the pressure exerted by neutralizing antibodies against hemagglutinin.

## Introduction

The influenza A virus (IAV) is an important human respiratory virus with a broad host range and primary reservoir in wild aquatic birds. In recent years, avian IAV has shown an alarming increase in spread across the globe and several spillovers to other species including domestic and wild-life mammals were detected resulting in high mortality rates [1–4]. To establish an infection in the human respiratory tract, IAV spreads through contact and aerosol and must overcome several pulmonary barriers, such as mucus and surfactant layer [5,6]. Morphologically, IAV is highly variable, ranging from spherical viruses of 100 nm in diameter to filamentous particles of more than 20 µm length [7]. Filamentous IAV is consistently found in human and animal isolates. This was also shown by electron microscopy (EM) of viral isolates from the last H1N1 pandemic in 2009 [8,9] and from H5N1 avian viruses [10]. In contrast, spherical particles are more prevalent in lab-adapted strains. Serial passage of clinical isolates in embryonated chicken eggs or Madine-Darby canine kidney (MDCK) cells frequently leads to a loss of filamentous morphology. Conversely, serial infection with spherical IAV in Guinea pigs has been shown to results in the emergence of filamentous virus particles [11].

IAV morphology is genetically dictated by the M segment [12], which is required for virus-like particle assembly [13]. This segment encodes for the M1 protein which forms a helical layer underneath the viral envelope [14], and for the M2 ion channel implicated in membrane scission [15]. Previous studies showed that the formation of filamentous virions depends on the actin cytoskeleton and Rab11-positive sorting endosomes [16,17,12]. Although it has been shown that a single mutation in the M segment can lead to a morphological switch from spherical to filamentous and *vice versa* [18], IAV morphology is not solely encoded in the genome. The high morphological variety within a single IAV clone was proposed to be driven by the tuneable assembly process in response to environmental pressure [19]. In addition, even in the scarcity of M1 and M2 proteins, the morphology of virions is maintained [20]. Regardless of morphology, infectious IAV incorporates only one set of a genome physically separated into 8 viral ribonucleoprotein complexes (vRNPs), positioned in the leading end of budding virions [21,22]. The assembly of long filamentous viruses presumably requires more time and larger amounts of hemagglutinin (HA) and matrix protein 1 (M1). As previously shown, NP:M1/HA ratios in filamentous particles are significantly lower than in spherical particles [12]. Laboratory-adapted spherical viruses are assumed to minimize the number of their structural proteins to encapsulate 8 viral ribonucleoprotein segments (vRNPs) during adaptation to cell culture.

The filamentous morphology has the advantage of providing a larger surface containing significantly more HA proteins required for entry. It is known that the IAV morphology dictates the route of entry. Filamentous virions predominantly enter by macropinocytosis, while spherical virions enter via clathrin-mediated endocytosis (CME) [23–25]. From the standpoint of viral fitness, high morphological variability within a virus population may therefore positively contribute to virus entry by exploiting more entry routes. Previous EM studies showed that small filamentous virions remain intact during cell entry. However, *in vitro* data revealed that filamentous virions disintegrate into spherical particles at endosomal pH [23], indicating that large filaments undergo more complex uncoating.

Previous studies demonstrated that filamentous morphology is important for IAV spread and transmissibility. Replacement of the M segment of the spherical, non-transmissive A/Puerto Rico/8/34 (H1N1) virus with that of the pandemic, filamentous IAV isolate A/Netherlands/602/2009 (H1N1) yields a virus with filamentous morphology which has indistinguishable transmissibility to wild type A/Netherlands/602/2009, as shown in a guinea pig transmission model [26].

Overall, the importance of IAV morphological heterogeneity is often overlooked in both *in vitro* and *in vivo* IAV studies. The current model states that the filamentous morphology increases virus fitness at higher cell entry pressure [19]. However, it is not fully understood why viruses have adapted to reduce morphological variability and minimize their shape to spheres in cell culture. In addition, the exact components of the pulmonary barriers which can be overcome by high morphological variability and filamentous morphology in physiological settings remain unidentified.

In our study, we generated reporter viruses encoding polymerase acidic proteins (PA) tagged with mScarlet that carry identical HA and neuraminidase (NA) and thus display identical antigenic surfaces but have a distinct spherical or filamentous morphology. We established a correlative fluorescent and scanning electron microscopy workflow to monitor the spread and morphology of virions at defined pulmonary or immunological pressures. We show that IAV infection in Calu-3 cells induces cell motility. Furthermore, we discovered that spherical viruses exhibit increased entry kinetics and spread faster in diverse tissue cultures, even in the presence of mucin or at variable cell densities, demonstrated in adhesion junction deficient cells. Strikingly, neutralizing antibodies against hemagglutinin are more effectively blocking virions with spherical morphology.

## Results

### Generation and characterization of IAV reporter viruses with predominant spherical or filamentous morphology

To exclusively compare IAV morphology-dependent effects, we generated two reporter viruses with distinct phenotypes, using a reverse genetics (RG) system [27] with genetically modified plasmids in an influenza A/WSN/33 (H1N1) (WSN) background. We used a plasmid encoding mScarlet, a fluorescent protein, fused to the PA gene to generate WSN:PAmScarlet (Figure 1A), based on previously published work [28]. The mScarlet gene was codon-optimized to remove CpG dinucleotides to evade recognition by Zinc Finger Antiviral Protein [29] and thereby increase the stability of the reporter viruses. As WSN is a lab-adapted strain with spherical morphology, we exchanged the WSN-M1 gene segment with the M1 segment of A/Udorn/307/72 (H3N2) to produce WSN-M1_Udorn_:PAmScarlet with a predominantly filamentous morphology (Figure 1 B, E), as previously published [20,19]. M1_Udorn_ contains 5 point mutations when compared to M1 from WSN (Figure 1 C). Cryo-electron microscopy (cryo-EM) analysis of viruses confirmed that WSN-M1_Udorn_:PAmScarlet contained 79.34% (n=219) of filamentous particles with a virion axis ratio > 2 and a median particle length of 547 nm. WSN:PAmScarlet viruses were mainly spherical (80.50%, n=70) (Figure 1F). The median length of WSN:PAmScarlet virions was 132 nm. Overall, 34.25% of WSN-M1_Udorn_ virions were longer than 1 µm compared to 2.86% for WSN. The maximum observed virion lengths were 4.78 µm for WSN-M1_Udorn_ and 1.51 µm for WSN. Interestingly, cryo-electron tomography (cryo-ET) of spherical virions revealed gaps and kinking of the M1 layer, presumably to accommodate vRNPs (Figure 1 D, arrowhead) during budding [30]. To validate that the reporter viruses carried the PA:mScarlet gene, we used fluorescence microscopy to image plaques formed in MDCK cells infected with WSN-M1_Udorn_:PAmScarlet or WSN:PAmScarlet (Suppl. Figure 1 A, B). This showed that 71% of WSN-M1_Udorn_:PAmScarlet and 42% of WSN:PAmScarlet rescued viruses expressed mScarlet (Suppl. Figure 1 C). The remaining fraction (29% and 58%) of viruses presumably eliminated the mScarlet. Note that non-fluorescent plaques were larger than fluorescent ones at 18 and 36 hpi, which indicates that mScarlet fused to PA yields a disadvantage to virus replication (Suppl. Figure 1 E).

**Figure 1:**
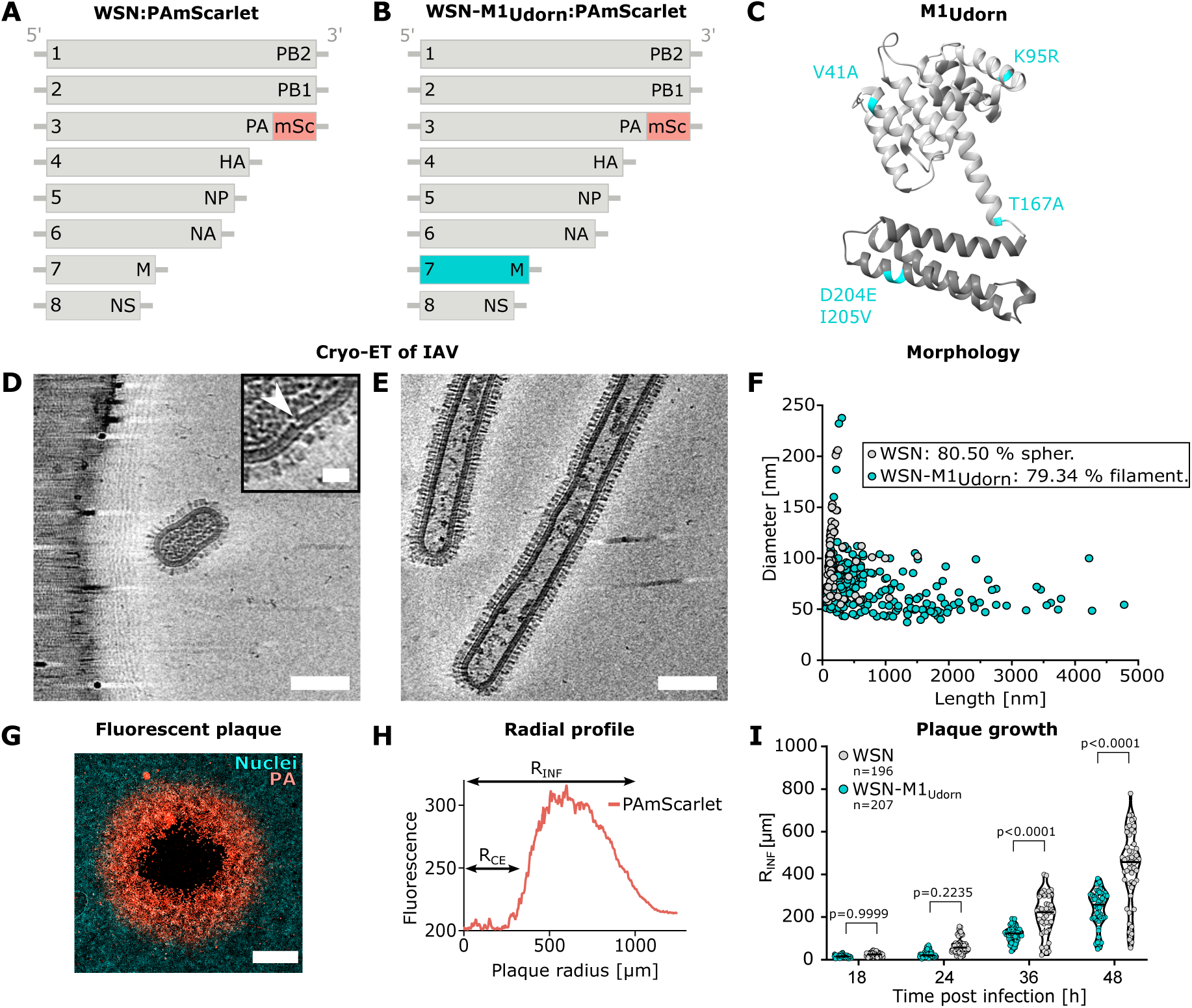
Morphological characteristics and spread of spherical and filamentous reporter IAVs. **(A)** Schematic representation of the eight genomic segments of A/WSN/33:PAmScarlet (WSN:mScarlet) with the fluorescent reporter mScarlet (mSc, red) fused to the PA gene segment. **(B)** Genomic segments of WSN:PAmScarlet, where the segment 7 was exchanged by the M segment of A/Udorn/72 (cyan) (WSN-M1_Udorn_:PAmScarlet). **(C)** Ribbon structure of the matrix protein 1 (M1) from A/WSN/33 (PDB: 6Z5L) harboring 5 amino acid substitutions (cyan) from M1 of A/Udorn/72. **(D)** Computed slices through a cryo-electron tomogram of an isolated WSN particle. A gap in the M1 layer is indicated by a white arrowhead. Scale bar of zoom-in 20 nm. **(E)** Slice through cryo-electron tomogram of two isolated filamentous WSN-M1_Udorn_ virions. Scale bars 100 nm. **(F)** Quantification of virion diameters and lengths for WSN (grey) (n=70) and WSN-M1_Udorn_ (cyan) (n=219) from TEM overview maps. Percentages of spherical (spher.: length/diameter ratio ≤ 2) and filamentous (filament.: length/diameter ratio > 2) phenotypes are indicated. **(G)** Exemplary fluorescence image of a plaque in a MDCK cell monolayer (cyan), initiated by the infection with a single virus particle and spread of released viruses from the center outward. The plaque can be divided into three zones from the center: empty zone caused by cytopathic effect (black), surrounded by an infected cell zone showing WSN:PAmScarlet signal (red), and uninfected cells (cyan). Scale bar 500 µm. **(H)** Quantification of viral growth dynamics by radial profile analysis of the cytopathic effect radius (R_CE_) from plaque center to the edge of empty zone and radius of infected cells (R_INF_) from plaque center to the outer edge of PAmScarlet positive cells. **(I)** R_INF_ of MDCK cells infected with WSN (grey) or WSN-M1_Udorn_ (cyan) at different time points (18, 24, 36, 48 h) post infection. The plaque growth for WSN and WSN-M1_Udorn_ infected cells were compared by ANOVA (p<0.0001) followed by Šídák’s multiple comparisons test. P-values are indicated.

### Quantification of fluorescent plaques demonstrates delayed spread of filamentous IAV

To quantitatively compare the spread of WSN-M1_Udorn_:PAmScarlet (filamentous) or WSN:PAmScarlet (spherical) viruses we analyzed each plaque using radial profile averaging of mScarlet fluorescence. We defined 3 different zones of infection representing detached cells (cytopathic effect), infected cells showing PAmScarlet signal, and uninfected cells (Figure 1 G and Suppl. Figure 1 D). The radial average of mScarlet signal showed a peak which allowed us to determine the radius of the cytopathic effect (R_CE_) and the infection radius (R_INF_) (Figure 1H). The analysis of 408 plaques revealed that spherical viruses spread faster than filamentous viruses in MDCK cells (Figure 1 I).

### Virus morphology does not change throughout the course of infection

The assembly of filamentous virions requires a larger number of structural proteins and possibly also takes a longer time before the particle is released to the media. This prompted us to analyze the morphology of budding virions during the infection and in particular to address whether filamentous morphology is lost in the course of infection within a plaque. We established a correlative fluorescence and scanning electron microscopy (CLSEM) method that can be applied to image virion morphology at different stages of infection spread within individual plaques (Figure 2 A, B). This correlative approach allowed us to analyze the morphology of virions at the surface of infected cells at 36 and 42 hpi when multiple rounds of infection had occurred. At the surface of MDCK cells infected with WSN:PAmScarlet, we observed a high number of spherical particles in proximity of the plaque center (Figure 2 E, yellow arrowhead), as well as at the border of the plaque (Figure 2 F, yellow arrowhead). On cells infected with WSN-M1_Udorn_:PAmScarlet, filamentous viruses of several micrometers in length were observed (Figure 2 K, L, cyan arrowheads). Remarkably, the morphology of filamentous and spherical virions was retained at all areas of the plaque (Figure 2). Uninfected cells showed numerous filopodia which have a wider and more variable diameter (69 ± 52 nm, n=10) than filamentous viruses. The data indicate that filamentous morphology is stable throughout multiple rounds of infection.

**Figure 2:**
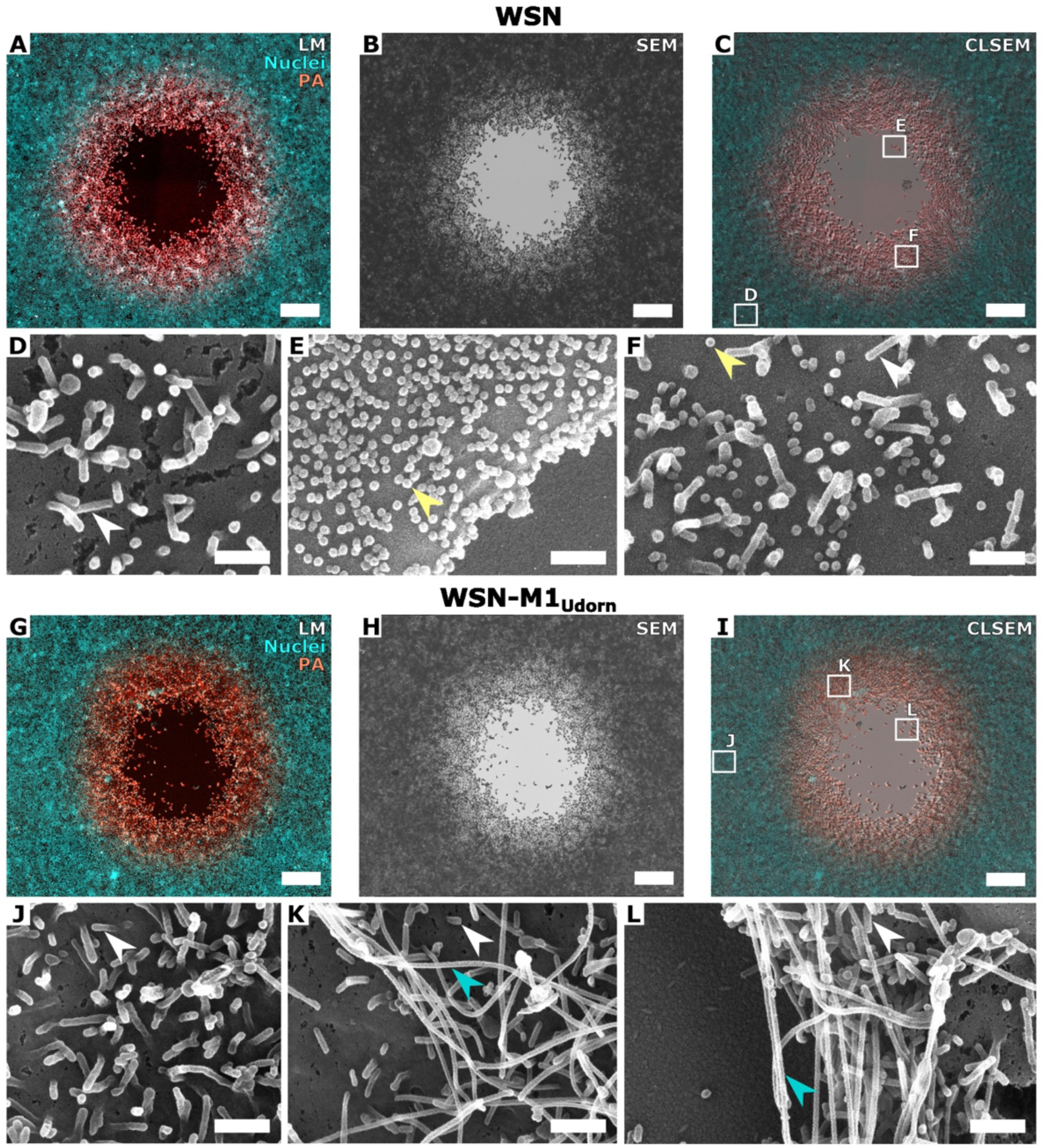
Correlative light and scanning electron microscopy (CLSEM) of IAV cell-to-cell spread. MDCK cells infected with spherical WSN:PA-mScarlet at 36 hpi (upper panel) or filamentous WSN-M1_Udorn_:PA-mScarlet at 42 hpi (lower panel). **(A, G)** Light microscopy images of fluorescent plaques showing PA-mScarlet (red) and cell nuclei (cyan). **(B, H)** Scanning electron microscopy images of plaques shown in A and G. **(C, I)** CLSEM of IAV-plaques. Scale bars 100 µm. **(D, J)** High-magnification SEM images of uninfected MDCK cell zones, **(E, K)** infected MDCK cells in the plaque center, **(F, L)** infected MDCK cells in the plaque periphery. Scale bars 0.5 µm. Yellow arrowheads show spherical viruses. Cyan arrowheads highlight filamentous particles. White arrowheads show filopodia.

### Filamentous IAV exhibit delayed entry kinetics

Since our data show that the filamentous phenotype confers a disadvantage in cell-to-cell spread in cell culture, we next assessed the entry kinetics of both viruses using an entry uptake assay in A549 cells. The duration of viral entry can be determined by inhibiting endosomal acidification with NH_4_Cl at various time points. Interestingly, this unveiled that the uptake of filamentous viruses is considerably delayed with an entry half-time of 33 min in contrast to the spherical viruses whose uptake was at least twice as fast (Figure 3 A). In addition, we could show that filamentous viruses at the same multiplicity of infection (MOI = 3) led to the infection of around 52% of cells, while spherical viruses could infect about 84% of cells (Figure 3 B). Inhibitors dynasore and 5-(N-Ethyl-N-isopropyl)amilorid (EIPA), which target CME and macropinocytosis, respectively, effectively blocked the infection by both spherical and filamentous viruses (Figure 3 C, D). However, morphology-selective inhibition was not detected. Overall, our data showed that spread and entry kinetics of filamentous viruses are reduced, indicating that the spherical morphology is better adapted to cell culture systems.

**Figure 3.**
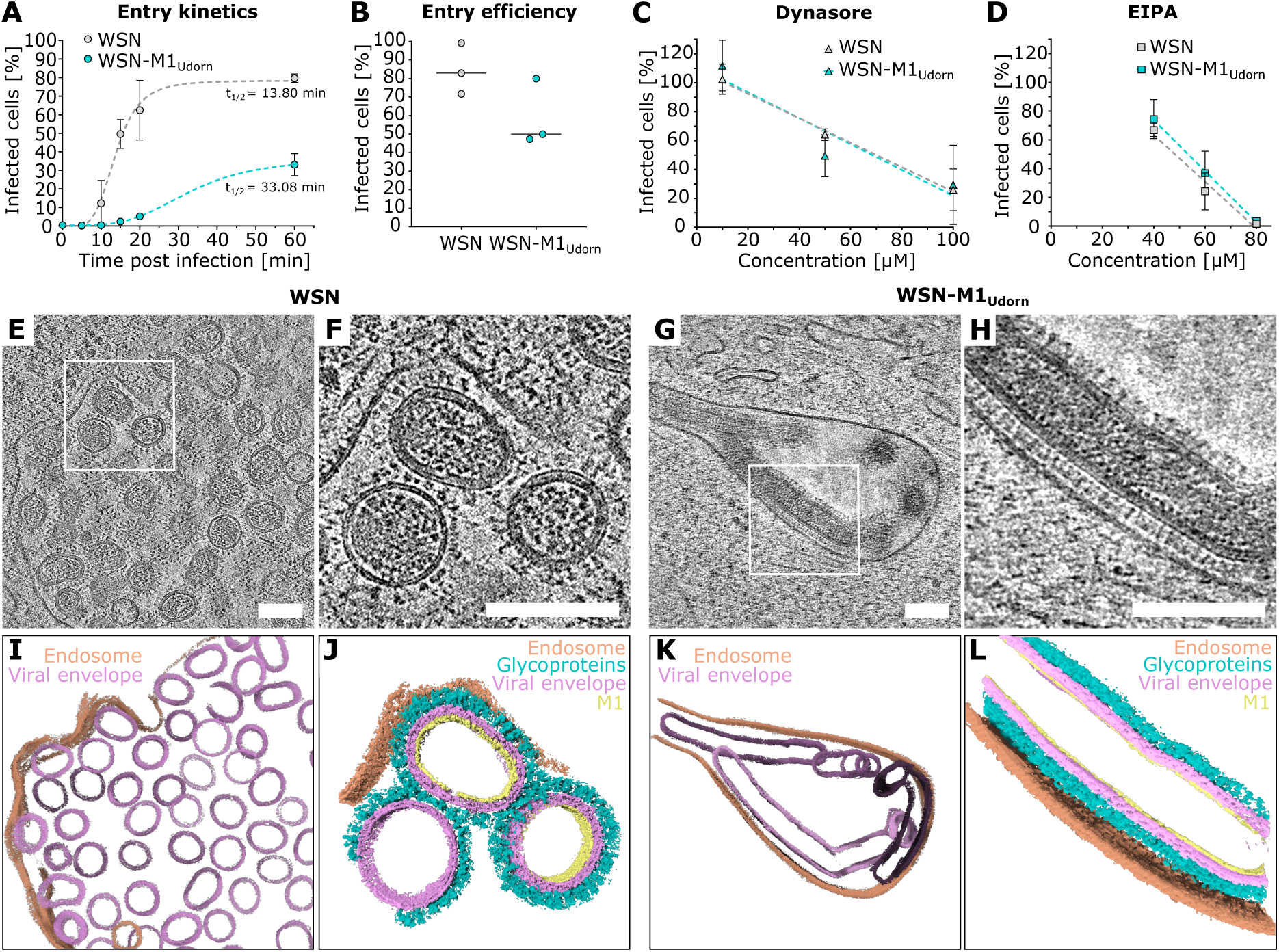
Temporal and structural analysis of spherical and filamentous IAV cell entry. **(A)** Entry time course of WSN (grey) and WSN-M1_Udorn_ (cyan) assessed by infection of A549 cells (MOI = 3) and treatment with NH_4_Cl (50 mM) at different time points (0, 5, 10, 15, 20, 60 min) post infection and fixation at 12 h after the last time point. Infected cells were quantified by fluorescence microscopy and a penetration half-time was determined based on a four-parameter logistic (4PL) curve in three independent experiments for each virus strain: R^2^(WSN)=0.943, t_1/2_(WSN)=13.80 min, R^2^(WSN-M1_Udorn_)=0.969, t_1/2_(WSN-M1_Udorn_)=33.08 min. **(B)** Percentage of infected A549 cells by WSN (grey) WSN WSN-M1_Udorn_ (cyan) (MOI=3, respectively) at 12 hpi determined by fluorescence microscopy in three independent experiments. Median(WSN)=83.72, median(WSN-M1_Udorn_)=51.86. **(C, D)** Comparison of inhibitory effect of increasing dynasore and EIPA concentrations on infection of A549 cells by WSN (grey) or WSN-M1_Udorn_ (cyan) (MOI=3 or 10), determined by fluorescence microscopy at 6 hpi. Cells were pre-treated with indicated drug concentrations for 1h prior to infection. Values were normalized to DMSO-treated control. Mean and standard deviations are indicated for three independent experiments. Linear fits for dynasore: R^2^(WSN)=0.925, R^2^(WSN-M1_Udorn_)=0.717 and EIPA: R^2^(WSN)=0.918, R^2^(WSN-M1_Udorn_)=0.899. Slices through cryo-electron tomograms (upper panel) showing **(E, F)** spherical WSN particles and **(G, H)** filamentous WSN-M1_Udorn_ particles in endosomal compartments of infected A549 cells at 15-30 min post infection and corresponding 3D segmentations **(I-L)**. Color code: endosomes (orange), viral envelope (pink), viral glycoproteins (turquoise), matrix protein 1 (M1) (yellow).

### Cryo-ET reveals intact filamentous viruses inside endosomal compartments

Given the delayed entry of filamentous IAV, we sought to visualize viral uptake and structurally characterize spherical and filamentous virions inside late endosomes using cellular cryo-ET. To this end, A549 cells were grown on EM grids and infected using a synchronized infection as done in our previous study [31]. Infected cells were vitrified and milled using a focused ion beam milling to generate electron-transparent lamellae with a thickness of about 200 nm. Consistent with the entry kinetics assay showing the effective uptake of spherical viruses, cryo-ET analysis of cells infected with the spherical virus showed a high number of virions inside the endosomal compartment (Figure 3 E, I). Interestingly, spherical virions showed distinct axis ratios based on the presence of the M1 layer (Suppl. Figure 3 D, E). While virions with an intact M1 layer showed an ovoidal shape with an axis ratio above 1, virions with a disassembled M1 layer were spherical (Suppl. Figure 3 E). This indicates that upon acidification, M1 layer disassembly leads to shape relaxation to a more spherical shape. We were able to find one endosome containing three long filamentous viruses (length > 500 nm) within a cell infected with WSN-M1_Udorn_ (Figure 3 G, H). Although 3D segmentation of the filamentous virions showed that they were bent inside the endosome, the particles were intact (Figure 3 K, L).

### IAV cell-to-cell spread in the absence of cell adhesion junctions and in the presence of mucin

Since filamentous viruses can be several micrometers long, we hypothesized that they could confer an advantage in cell-to-cell spread when the contact between cells is disrupted. To test this hypothesis, we used MDCK-α-Catenin knock-out (KO) cells [32] that show deficient adherens junctions and do not form a monolayer (Figure 4 A). This is reflected by a decreased cell density of MDCK-α-Catenin-KO cells (5821 cells/0.15 cm^2^) as compared to MDCK cells (7627 cells/0.15 cm^2^) (Figure 4 B). However, even at lower cell densities, IAV with spherical morphology has an increased cell-to-cell spread compared to the filamentous virus (Figure 4 C). We next tested mucin, which represents an important component of the mucus layer and is a pulmonary barrier known to inhibit IAV infection and can be penetrated using NA activity (Figure 4 D) [33–35]. To evaluate the inhibitory effect, mucin was mixed into an agarose overlay at a concentration range of 0.5-2% and analyzed by fluorescent plaque assays in MDCK cells. Since the inhibitory effect of mucin could also be due to a change in the viscosity of the agarose gel, we first evaluated the physical properties of the mucin-agarose mixtures. Our data revealed that the viscosity and elasticity of the mixture does not strongly depend on mucin concentrations (Figure 4 E). Subsequently, we investigated the impact of mucin on IAV cell-to-cell spread. Radial profile averaging of fluorescent plaques revealed that mucin effectively inhibits IAV spread. However, mucin did not show any IAV morphology-dependent inhibitory effect as indicated by the slopes of linear fits of the plaque diameters for WSN:PAmScarlet and WSN-M1_Udorn_:PAmScarlet (Figure 4 F). To evaluate the cell-to-cell spread of spherical and filamentous IAV in lung cells, we next performed fluorescent plaque assays in Calu-3 cells. As opposed to MDCK cells, Calu-3 cells are bronchial epithelial cells naturally producing mucins [36]. Interestingly, infection in Calu-3 cells did not result in plaques but in infection foci (Figure 4 G). Furthermore, our data show that filamentous viruses spread slower than spherical viruses also in Calu-3 cells. Time-lapse imaging of Calu-3 foci revealed that cells infected by either WSN:PAmScarlet or WSN-M1_Udorn_:PAmScarlet migrate towards the center of the foci (Suppl. Figure 4 A, E). The cell migration velocity, directionality and distance migrated towards the focus center were increased in IAV-infected Calu-3 cells as compared to uninfected cells (Suppl. Figure 4 B-F).

**Figure 4:**
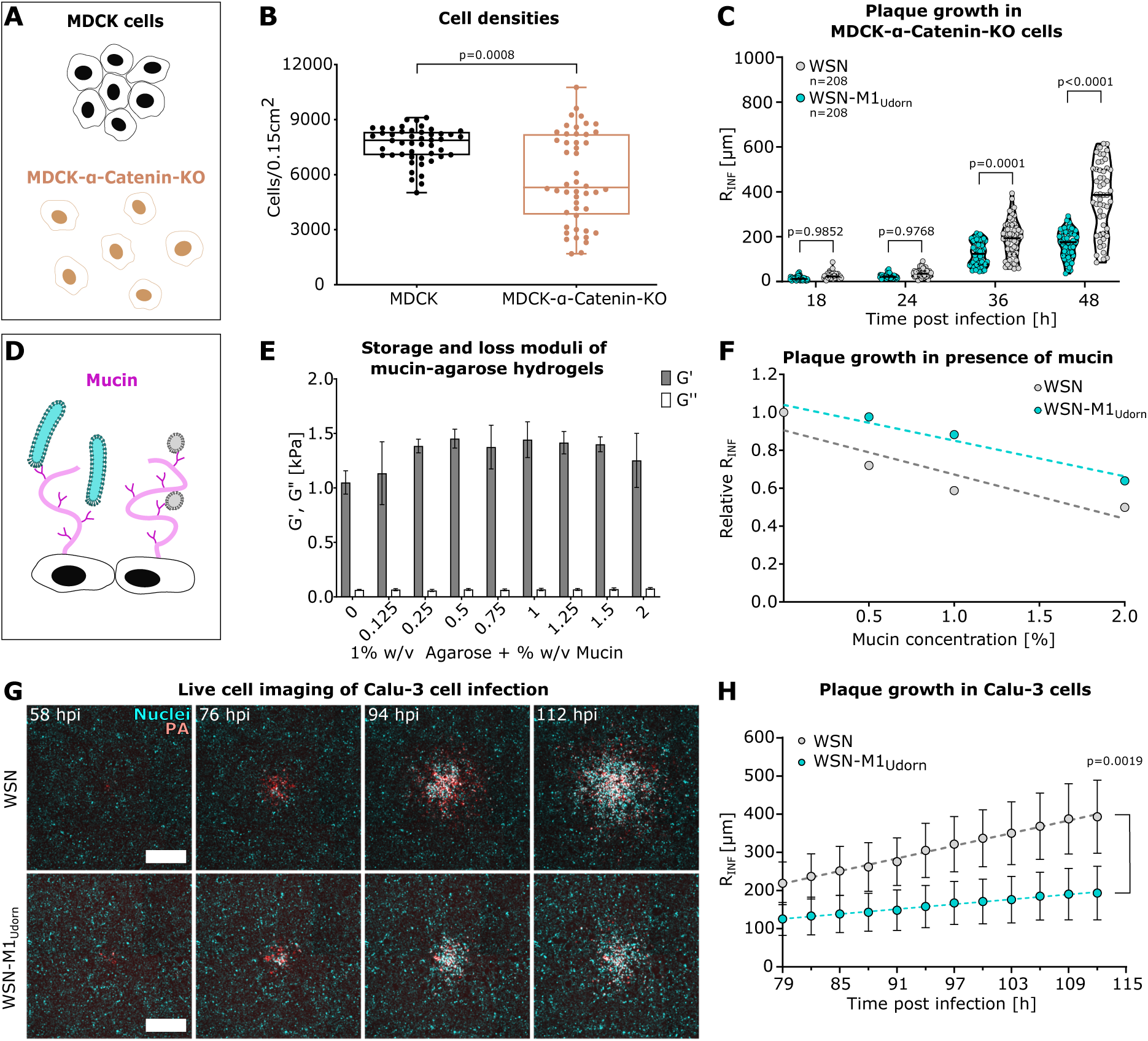
Impact of cell-cell contact and mucin on IAV spread. **(A)** Schematic diagram illustrating that MDCK cells form a uniform monolayer. In MDCK-α-Catenin-KO cells are impaired in forming a monolayer, due to disrupted adherens junctions. **(B)** Number of MDCK cells (black) and MDCK-α-Catenin-KO cells (brown) per area of 0.15 cm^2^. Data points show cell counts from 48 positions of 3 replicates. Boxes represent mean and interquartile ranges. Whiskers show the data range. The p-value was determined by two-tailed t-test. **(C)** R_INF_ of MDCK-α-Catenin-KO cells infected with WSN (grey) or WSN-M1_Udorn_ (cyan) at different time points (18, 24, 36, 48 h) post infection. For each time point, the plaque growth for WSN and WSN-M1_Udorn_ infected cells were compared by ANOVA (p<0.0001) followed by Tukey’s multiple comparisons test. P-values are indicated. **(D)** Schematic structure of mucins, highly glycosylated proteins and components of the mucosal barrier of the lung epithelium. Sialic acids displayed on mucins can capture IAV via binding to hemagglutinin. The sialidase activity of NA mediates mucus penetration of IAV. **(E)** Rheological properties of agarose hydrogels in presence of increasing mucin concentrations. The storage modulus (G’, grey) reflects elasticity and the loss modulus (G’’, white) represents viscosity. Bars show averages of 3-4 repeated measurements in 5 different mucin-agarose hydrogels with indicated standard deviations. **(F)** Relative R_INF_ of WSN (grey) and WSN-M1_Udorn_ (cyan) in presence of increasing mucin concentrations, normalized to untreated cells. Linear fits are shown: R^2^(WSN-M1_Udorn_) = 0.829, R^2^(WSN-M1_Udorn_) = 0.949**. (G)** Exemplary fluorescence microscopy images of WSN and WSN-M1_Udorn_ plaques from live cell imaging of Calu-3 cells at indicated time points with cell nuclei shown in cyan and IAV-PA in red. Scale bars: 500 µm. **(H)** R_INF_ determined by live cell imaging of Calu-3 cells infected with WSN (grey) or WSN-M1_Udorn_ (cyan) between 79 and 112 hpi. Each data point represents the mean of at least 16 plaques from 2 independent experiments. Standard deviations are indicated. Linear fits are indicated: R^2^(WSN-M1_Udorn_) = 0.994, R^2^(WSN-M1_Udorn_) = 0.993. The p-value for 112hpi was determined by Mann Whitney test.

### IAV cell-to-cell spread in the presence of hemagglutinin-binding neutralizing antibody

Filamentous virions contain significantly more hemagglutinin glycoproteins than spherical virions. Based on the length distribution (Fig 1) and HA-HA spacing [37], we estimated that filamentous virions have on average 3.5 times more HAs (Figure 5 A). This prompted us to investigate whether filamentous virions will be able to escape from neutralization by a stalk-binding antibody MEDI8852, which has been shown to limit transmission of pandemic IAV [38]. While we could observe that neutralizing antibodies inhibit the spread of viruses, a higher concentration of antibodies was needed to inhibit filamentous viruses (IC50 = 1.6 nM) compared to spherical viruses (IC50 = 0.9 nM) (Figure 5 B and 5 C). This was further confirmed by correlative fluorescence and CLSEM imaging of plaques. In the presence of the neutralizing antibodies, we observed a significantly lower number of spherical WSN:PAmScarlet virions on the cell surface as compared to untreated infected cells (Figure 5 D-F). In contrast, the number of filamentous virions on the cell surface infected with WSN-M1_Udorn_:PAmScarlet did not change significantly upon antibody treatment (Figure 5 G-I).

**Figure 5:**
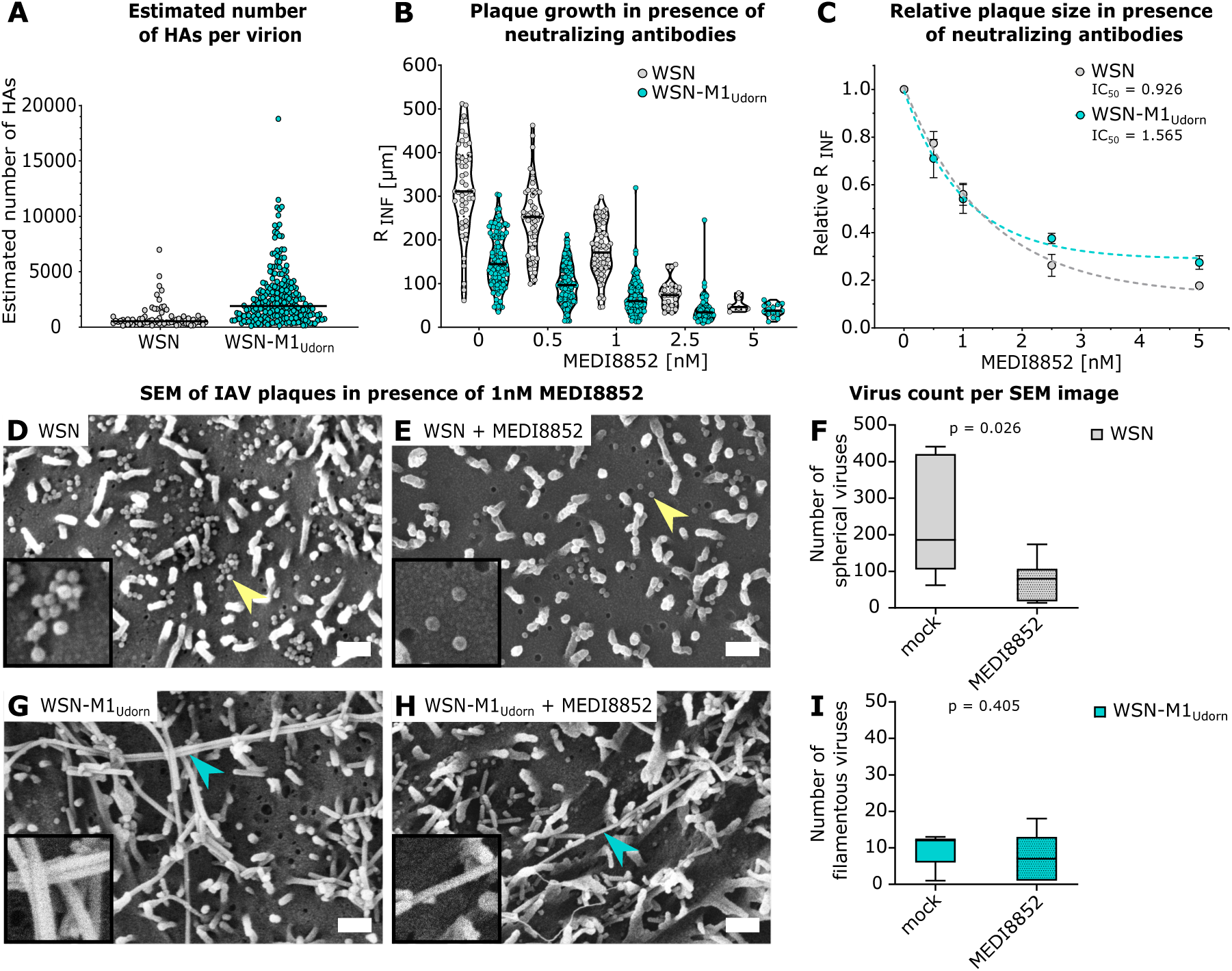
Effect of neutralizing anti-HA antibodies on morphology-dependent IAV spread. **(A)** Estimated number of HAs per virion surface based on size measurements from Fig. 1F. Median(WSN) = 534.73, median(WSN-M1_Udorn_) = 1904.31. **(B)** Infection radius (R_INF_) in MDCK cells infected with WSN (grey, n = 388) or WSN-M1_Udorn_ (cyan, n = 921) in presence of increasing concentrations of MEDI8852 at 36 hpi. **(C)** Relative R_INF_ for WSN and WSN-M1_Udorn_ in presence of 0.5, 1, 2.5 or 5 nM MEDI8852, normalized to untreated controls. Mean values from three independent experiments and standard deviations are indicated. IC50 was determined from three-parameter logistic (3PL) curve fits: R^2^(WSN) = 0.926, IC50(WSN) = 0.925 nM, R^2^(WSN-M1_Udorn_) = 0.974, IC50(WSN-M1_Udorn_) = 1.565 nM. **(D)** Scanning electron microscopy (SEM) images of plaques in MDCK cells infected with WSN without neutralizing antibody and **(E)** in presence of 1 nM MEDI8852. Yellow arrowheads indicate spherical viruses. Scale bars 3 µm. **(F)** Number of spherical viruses per image for mock treated (median = 174.5) and antibody treated (1 nM MEDI8852, median = 80.0) MDCK cells from 7 plaques. **(G)** SEM images of plaques in MDCK cells infected with WSN-M1_Udorn_ without neutralizing antibody and **(H)** in presence of 1 nM MEDI8852. Cyan arrowheads indicate filamentous viruses. Scale bars 3 µm. **(I)** Number of filamentous viruses per image for mock treated (median = 12) and antibody treated (1 nM MEDI8852, median = 7) MDCK cells from 5 plaques.

## Discussion

While the pleomorphic nature of IAV particles directly contributes to higher infectivity and transmissibility in vivo, spherical morphology is selected for in cell culture [11]. Recent studies have characterized heterogeneous IAV particles on a structural [7,39,21] and genetic level and identified residues in the M1 protein as essential determinants of virion morphology [20,40]. However, the functional role of filamentous virions in the infected host and factors favoring this phenotype remain to be identified. One reason for this gap is the lack of quantitative approaches that systematically compare different virion morphologies in various conditions. Here we investigated morphology-linked differences in host cell entry and cell-to-cell spread of IAV in vitro. By establishing a fluorescent reporter virus systems of distinct spherical and filamentous phenotypes as described by Bourmakina, Garcia-Sastre [20], Vahey and Fletcher [41] but identical surface antigens, we were able to track infection spread over time. Notably, the rescued reporter viruses used in this study exhibit around 80% of the phenotype. These results are in line with previous studies showing that M1 is the determinant for IAV morphology [21,42]. Our data demonstrate a disadvantage of filamentous virions in cell-to-cell spread in MDCK cells and an MDCK-α-Catenin knock-out cell line which shows increased cell-to-cell distance in the monolayer. Hence, our data indicate that filamentous viruses do not gain an advantage in cell-to-cell spread kinetics at low cell densities as we initially hypothesized.

The delay in the spread of filamentous viruses can occur at different stages of viral replication, and likely entry is one of the major factors. Since both viruses have the same genetic background the differences in virus replication cycle kinetics likely occur on entry or exit levels. We showed that the endosomal escape of filamentous viruses is delayed by 20 min compared to spherical viruses. Delayed early infection of filamentous viruses was also reported in a study using spherical A/Udorn/307/72 and filamentous A/Udorn/307/72 10A variants [43]. Consistent with existing evidence showing that the entry of both filamentous and spherical IAV is blocked by inhibitors targeting macropinocytosis [23], our data show that the entry of spherical and filamentous viruses is inhibited by EIPA or by dynasore to a similar extent. Since EIPA inhibits the acidification of endosomes [23,44], entry inhibition of spherical viruses cannot be excluded. Moreover, it has been shown previously that both spherical and filamentous viruses can induce macropinocytosis [24,23]. Hence, both spherical and filamentous viruses are likely to utilize multiple entry pathways.

It is well-established that filamentous viruses undergo disintegration upon low pH treatment in vitro [23]. Hence, we analyzed the structure of spherical and filamentous virions inside the endosomes by cellular cryo-ET of cryo-focused ion beam-milled infected cells. This allowed us to capture long filamentous virions inside endosomes, which were bent but did not undergo disintegration into smaller components and were longer than those reported previously [23]. However, this disintegration is pH dependent and long filamentous virions were most likely in early endosomes as the hemagglutinin was in a prefusion conformation. Notably, the endosomal environment led to morphological changes of spherical virions, which upon M1 layer disassembly became rounder.

Hence, our findings indicate that the increased entry kinetics of spherical viruses is a factor which drives the morphological adaptation towards spherical virions in vitro. However, other factors likely contribute to this adaptation, such as increased assembly efficiency for spherical viruses which require fewer building blocks for assembly. Previous studies using video-microscopy imaging of the budding respiratory syncytial virus revealed an average speed of filament elongation of 110-250 nm/s [45]. Assuming a similar budding velocity in IAV, it is unlikely that budding and growth of long filamentous virions significantly contributes to the delay in the cell-to-cell spread of filamentous viruses. The accumulation of spherical virions observed by SEM on the cell surface indicates that the rate-limited step in budding is likely membrane scission rather than budding particle growth.

Since filamentous IAV spread is slower in cell culture, we hypothesized that they undergo morphological adaptation towards a spherical morphology throughout infection. In order to assess whether the filamentous morphology is lost during several rounds of infection in cell culture [11], we established a CLSEM approach to image budding virions within different zones of a plaque using SEM. Our results show that the morphology of budding virions is not altered as infection progresses. This indicates that virus clone propagation within a single plaque might not undergo sufficient replication cycles for morphology adaptation to occur. Alternatively, morphology maintenance or changes driven by adaptation may require a selective pressure that is lacking in vitro.

To investigate different types of selective pressure, we examined the role of mucin and broadly neutralizing anti-HA stalk antibodies. Mucins limit IAV infection in vivo [46] and are known to be induced upon interferon treatment or IAV infection, and to effectively reduce the infection of different influenza A viruses [47]. Our data confirmed that the addition of mucin into the agarose overlay inhibits IAV infection in a dose-dependent manner. However, neither IAV with filamentous nor spherical morphology appear to have an advantage when compared to a mucin-free control. Similar results were obtained using Calu-3 cells, which are known to produce mucin. Interestingly, IAV with spherical morphology showed increased cell-to-cell spread also in Calu-3 cells. Unlike in MDCK cells, the infection in Calu-3 cells resulted in the formation of infection foci rather than plaque. Surprisingly, time-lapse movies revealed that infected Calu-3 cells move faster than uninfected cells towards the center of the foci. While this needs to be further studied, the data indicate that Calu-3 cells are migrating towards the center to replace the dying cells. In addition, these results suggest that IAV infection triggers cell motility, which was reported for example in vaccinia virus-infected cells [48].

Importantly, we could show that the presence of neutralizing anti-HA stalk antibodies leads to the loss of the cell-to-cell spread advantage of spherical viruses. This finding is in agreement with previous evidence showing that neutralizing antibodies have a stronger effect on spherical IAV [49], and suggests that filamentous viruses can better escape neutralization by antibodies due to a much higher number of hemagglutinins. Consistent with our results, a recent preprint showed that antibody pressure drives filament assembly [50].

Overall, our study suggests that morphology-dependent virus entry kinetics plays an important role in the cell-to-cell kinetics of IAV spread. Our data provide further evidence that IAV filamentous morphology is lost to accelerate cell-to-cell spread by faster entry kinetics and to achieve higher entry efficiency. However, this fitness penalty, which applies to filamentous viruses in cell culture, might be compensated in vivo due to neutralizing antibodies.

## Materials and Methods

### Cell lines

HEK 293T (human embryonic kidney 293T) cells were kindly provided by Dr. Marco Binder (DKFZ, Heidelberg, Germany). MDCK (Madin-Darby canine kidney) cells were a gift from Dr. Maria João Amorim (Instituto Gulbenkian de Ciência, Oeiras, Portugal). The cell lines were maintained in DMEM-GlutaMAX^TM^-I medium (ThermoFisher Scientific, Gibco) supplemented with 10% fetal bovine serum (FBS) (ThermoFisher Scientific, Gibco) and 1% penicillin/streptomycin (P/S) (ThermoFisher Scientific). Calu-3 (human lung cancer) cells, provided by Prof. Dr. Ralf Bartenschlager (Department of Infectious Diseases, Heidelberg University, Germany) were grown in DMEM-GlutaMAX^TM^-I medium supplemented with 20% FBS, 1% P/S and 10 mM sodium pyruvate (Sigma-Aldrich). A549 (human alveolar basal epithelial adenocarcinoma) cells [51] were obtained from ATCC and maintained in DMEM-F12 supplemented with GlutaMAX^TM^-I (ThermoFisher Scientific), 10% FBS and 1% P/S. All cells were cultured at 37°C in a humidified 5% CO_2_ atmosphere and passaged twice a week in a 1:10 ratio or 1:3 (for Calu-3 cells). Cell lines were tested for Mycoplasma contamination every 3 months.

### Fluorescent influenza A reporter viruses

All work with infectious IAV was performed under biosafety level (BSL-)2 conditions at BioQuant or CIID (Heidelberg University, Germany), following approved operating procedures. Viruses used in this study were rescued by reverse genetics (RG), using 8 bidirectional plasmids as described by [27]. Two reporter virus strains with the genetic background of influenza A/WSN/33, carrying PA tagged with a codon optimized mScarlet, were produced based on [52,28,31]. Predominantly spherical WSN:PAmScarlet was rescued by transfecting the RG plasmids: pHW2000-PB2-WSN, pHW2000-PB1-WSN, pHW2000-PA-WSN-mScarlet, pHW2000-HA-WSN, pHW2000-NP-WSN, pHW2000-NA-WSN, pHW2000-M-WSN, pHW2000-NS-WSN. Plasmids were obtained from Dr. Ervin Fodor (University of Oxford, UK) and Dr. Andrew Mehle (University of Wisconsin, Madison, USA). WSN-M1_Udorn_:PAmScarlet was produced by replacing plasmid pHW2000-M-WSN with pcDNA3.1-M1-Udorn-M2-WSN, kindly provided by Dr. Michael Vahey (Washington University, St Louis, MO, USA) and Prof. Daniel Fletcher (University of California Berkeley, Berkeley, CA, USA). This plasmid contains the M1 sequence from influenza A/Udorn/72 with 5 amino acid substitutions as compared to M1 from A/WSN/33 and was previously shown to confer a predominantly filamentous virion phenotype [41].

HEK 293T cells were seeded into 10 cm cell culture dishes at a density of 4×10^6^ cells per dish and grown over night. The transfection mix was prepared by mixing 2 ml Opti-MEM medium with 2.5 µg of each of the 8 RG plasmids and 60 µl TransIT® -LT1 transfection reagent (Mirus Bio) and incubation at room temperature (RT) for 30 min. The mix was subsequently added dropwise to the cells followed by incubation for 24h at 37°C and 5% CO_2_. The cell culture medium was replaced by FBS-free DMEM-GlutaMAX^TM^-I supplemented with 1% P/S, 0.3% BSA (Sigma-Aldrich) and 2 µg/µl TPCK-trypsin (Sigma-Aldrich). 4×10^6^ MDCK cells were seeded on the transfected HEK 293T cells for co-culture. Once a cytopathic effect was visible, cell supernatant was harvested and cleared from cell debris by centrifugation at 1,000 g for 10 min. The virus-containing supernatant (P0) was snap-frozen in LN_2_ and stored at -80°C.

### Plaque assay for titer determination

MDCK cells were seeded into 6-well plates (Corning) at a density of 1×10^6^ cells/well and grown into a monolayer overnight. Plaque assays were performed in duplicates. IAV aliquots stored at -80°C were slowly thawed on ice for one hour. A virus dilution series from 10^-3^ to 10^-8^ was prepared in 4°C cold FBS-free DMEM-GlutaMAX^TM^-I medium. The cell monolayer was washed once with FBS-free DMEM-GlutaMAX^TM^-I medium to remove residual FBS. Next, 0.8 ml of the virus dilutions were added to the cells followed by incubation at 37°C and 5% CO_2_ to allow virus attachment and adsorption. Unbound virus was aspirated, and cells were washed with PBS (Sigma-Aldrich). Each well was overlayed with 3 ml microcrystalline cellulose overlay consisting of FBS-free 2x DMEM medium supplemented with 2% P/S, 50 mM HEPES (Carl Roth GmbH + Co. KG), 7.4 g/l NaHCO_3_ (Sigma-Aldrich), 0.3% BSA and 2 µg/µl TPCK-Trypsin that was mixed with 2.4% Avicel (FMC) at a 1:1 ratio. After incubation at 37°C and 5% CO_2_ for 48 h without moving the plate, cells were washed 3 times with PBS and fixed with 4%PFA in PBS for 30 min. Crystal violet (Sigma-Aldrich) staining (1% in H_2_0) was performed for 10 min at RT followed by 3 washing steps with H_2_0. Plaques were manually counted and the average virus titer (T_virus_) from all dilutions was determined by

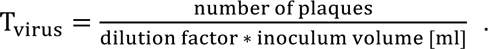

### Morphological characterization of viruses

To morphologically characterize the produced IAV strains by cryo-EM, virus-containing supernatants were thawed on ice for one hour. In the meantime, 200 mesh, cupper R2/1 grids (Quantifoil) were glow discharged. Undiluted virus was mixed with 10 nm protein A-coated colloidal gold (Aurion). 3 µl virus were applied onto the grids prior to plunge-freezing. Plunge-freezing into liquid ethane was performed using an automatic EM GP2 plunge-freezing device (Leica) under the following conditions: chamber temperature: 25°C, humidity: 80%, back-side blotting: 2-3 sec. Grids were stored in LN_2_ until imaging. Cryo-TEM data was collected with SerialEM [53] using a Titan Krios cryo-TEM (ThermoFisher Scientific) operated at 300 keV and equipped with a Quanta Imaging Filter (Gatan) with an energy filter slit set to 20 eV and a K3 direct electron detector (Gatan). First, grids were mapped at 8,700 x magnification (pixel spacing: 10.64 Å). From these maps, the length and diameter of virions from outer membrane to outer membrane was measured in IMOD [54]. For filamentous virions, the diameter was measured at 3 locations and averaged. Measurements were plotted Prism 10.1.2 (GraphPad). At representative positions of the map, tilt series were acquired at 33,000 x magnification (pixel spacing: 2.67 Å) using Parallel Cryo-Electron Tomography (PACEtomo) [55] in SerialEM [53] with the following setup: dose-symmetric tilting scheme [56], nominal tilt range from 60° to -60° and 3° increments, target defocus -3 µm, electron dose per record 3e^-^/Å^2^. Drift correction of acquired movies was done with Motioncor2 [57]. Tomograms were reconstructed in Etomo using the following parameters: tilt series alignment with patch tracking, weighted back projection with simultaneous iterative reconstruction technique (SIRT)-like filter equivalent to 5 iterations, dose-weighting, 3D contrast transfer function (CTF) correction.

### Fluorescent plaque assays

For fluorescent plaque assays, MDCK, MDCK-α-catenin-KO, or Calu-3 cells were seeded at a density of 1×10^6^ cells/well in 6-well plates and grown into a monolayer. IAV aliquots were thawed on ice for one hour and diluted to 50-100 PFU/well in FBS-free DMEM-GlutaMAX^TM^-I medium. Cells were washed once with FBS-free DMEM-GlutaMAX^TM^-I before infection for 1h at 37°C and 5% CO_2_. The monolayer was washed 2 times with PBS and overlayed with avicel overlay as described above. At 18, 24, 36 or 48 hpi, the overlay was removed, cells were washed 3 times with PBS and chemically fixed with 4% PFA (Electron Microscopy Science) in PBS for 30 min at RT.

For additional immunofluorescence staining of IAV-NP or M2, cells were washed three times with PBS and subsequently permeabilized with 0.2% Triton X-100 (Sigma-Aldrich) in PBS for 5 min at RT. Next, cells were washed 3 times with PBS for 5 min and blocking was performed for 30-60 min with 3% BSA in PBS-T (PBS supplemented with 0.1% Tween-20 (Roth)). After one washing step with dilution buffer (1% BSA in PBS-T), cells were incubated for 1h at RT with mouse anti-NP (MAB8257, Merck) or mouse anti-M2 (ab5416, Abcam) primary antibodies diluted 1:500 in dilution buffer. Cells were then washed 3 times for 5 min with PBS-T before adding the secondary goat anti-mouse Alexa Fluor^TM^ 488 antibody (A11029, ThermoFisher Scientific) and DAPI (Sigma-Aldrich), both at a dilution of 1:1000 in PBS-T. Plates were incubated for 1h at RT in the dark and subsequently washed 3 times with PBS.

Fluorescence microscopy data was acquired using the 5x objective lens of the Zeiss CellDiscoverer 7 microscope equipped with an Axiocam 712 camera. Tile sets covering 80% of the well were acquired in three channels with LED excitation of DAPI at 385 nm, Alexa Fluor^TM^ 488 at 470 nm and mScarlet at 657 nm. Tiles were stitched in the ZEN blue software (Zeiss).

### Analysis of IAV cell-to-cell spread

Cell-to-cell spread of spherical and filamentous reporter IAV was assessed by radial density profile measurements in ImageJ/Fiji [58,59]. For each plaque, a circle was drawn above the center of each plaque and the fluorescent signal of PA-mScarlet, NP or M2 was radially averaged. The cytopathic effect radius represents the distance from the center of the plaque and the inner edge of infected cells (Suppl. Figure 1 D). The infection radius (R_INF_) was defined as the distance between the center of the plaque and the outer edge of infected cells. These parameters were calculated for different time points after infection. Individual parameters and regression equations were plotted in Prism 10.1.2 (GraphPad). To assess the reporter efficiency, growth dynamics (R_INF_) of plaques resulting from spherical WSN and reporter WSN:PAmScarlet viruses were determined from M2 signals of WSN and PA-mScarlet signals of WSN:PAmScarlet for 10 plaques at 18, 24 and 36 hpi, respectively.

### Correlative light and scanning electron microscopy (CLSEM) of plaques

For SEM, MDCK cells were grown on indium tin oxide (ITO)-coated coverslips placed in 6-well plates until confluent and infected with fluorescent reporter IAV as described above (Suppl. Figure 2). For plaque assays in presence of neutralizing antibodies, 1nM MEDI8852 antibody was added to the overlay medium. After chemical fixation, DAPI staining was performed at a 1:1000 dilution for 5 min. Plaques were imaged using the 5x objective of the Zeiss CellDiscoverer 7 microscope. After fluorescent image acquisition, cover slips were washed with PHEM (15 mM PIPES, 6.25 mM HEPES, 2.5 mM EGTA, 0.5 mM MgCl_2_) or 0.1 M Cacodylate buffer. Next, cells were incubated with 1% OsO_4_ in cacodylate or PHEM buffer at 4°C for 30 min, followed by washing with Cacodylate or PHEM buffer (Suppl. Figure 2). Dehydration was achieved by addition of increasing concentrations of acetone (25%, 50%, 75%, 95%, 100%) to the sample and incubation for 10 min, respectively. Lastly, critical point drying was done on a Leica CPD300 (Leica Microsystems) at 17°C and 63.5 bar and the sample was sputter coated with a 5 nm thick layer of Au/Pd (80/20) using the Leica ACE600 (Leica Microsystems). Samples were mapped by SEM using an Aquilos 2 dual-beam cryo-focused ion beam-scanning electron microscope (ThermoFisher Scientific) at magnifications between 100 x and 50,000 x using 5 keV and 0.1 nA. Correlation with LM images was performed in the MAPS 3.3 software (ThermoFisher Scientific). The cell topology and virus morphologies were quantified with Fiji. For each condition, virus counts were determined from at least 4 images at 50,000 x magnification.

### Estimation of HA count per virion

To estimate the number of HA proteins per virion, the virion surface area was calculated based on particle length and diameter measurements from cryo-EM maps described above. For spherical IAV, we assumed perfect spherical symmetry using the formula A = π * d^2^. The basis for filamentous virion surface area is a cylinder with A = 2πr^2^ + 2πrl. Subsequently, the number of HAs per virion was calculated by assuming 7 proteins per 500 nm^2^ as described in [37].

### Cell densities

Cell densities of MDCK and MDCK-MDCK-α-catenin-KO cells were determined by segmentation of cell nuclei from uninfected cells using the StarDist plugin in ImageJ/FIJI [58,60]. Three areas of 0.15 cm^2^ were selected from independent experiments.

### Plaque assay in presence of mucins

Dose-dependent effects of mucins on IAV cell-to-cell spread were studied by plaque assay in presence of mucin from porcine stomach type III (M1778, Sigma-Aldrich) in a concentration range of 0-2% (w/v), consistent with physiological conditions [61] and technical limitations. A 5% (w/v) mucin stock solution was prepared by dissolving mucin in 2 x DMEM (Sigma-Aldrich) by stirring at 37°C for 1h. The stock was then UV-inactivated for 15 min to avoid microbial contamination and further diluted to 2%, 1% and 0.5% in 2 x DMEM. The mixtures were supplemented with 2% P/S, 50 mM HEPES, 0.6% BSA and 4 µg/µl TPCK-Trypsin. A confluent MDCK cell monolayer in 6-well plates was infected with a dilution of WSN:PAmScarlet or WSN-M1_Udorn_:PAmScarlet as described earlier. Meanwhile, mucin dilutions were mixed with 1% (w/v) agarose (Sigma-Aldrich) heated to 56°C. Infected cells were overlayed with the mucin-agarose hydrogel and incubated at 37°C and 5% CO_2_ for 60 h. Fluorescent plaques were analyzed according to the above-mentioned protocol.

### Physical properties of mucin-agarose hydrogels

Rheological properties of mucin-agarose hydrogels were performed using a HR 20 Discovery Hybrid Rheometer manufactured by TA Instruments^TM^. A frequency sweep with a fixed strain and a strain sweep with a fixed frequency were carried out to establish the optimal parameters. Based on these preliminary experiments, a strain of 0.25% and a frequency of 2 Hz were identified as suitable conditions for mucin-agarose hydrogels. The acquired experimental data were analyzed using the TRIOS software. Two key parameters, namely the storage modulus (G’) and the loss modulus (G’’), were measured. G’ reflects the material’s elasticity, providing insights into its ability to store and recover energy. G’’ represents the material’s viscosity, indicating its resistance to flow. To ensure the reliability of the results, each gel sample underwent 3 to 4 repeated measurements, and the obtained values were averaged. Results of G‘ and G‘‘ were plotted in Prism 10.1.2 (GraphPad). Error bars show standard deviations.

### Plaque assay in presence of neutralizing anti-HA antibodies (MEDI8852)

The effect of anti-HA broadly neutralizing antibodies (MEDI8852) [62] on morphology-dependent IAV cell-to-cell spread was tested by fluorescent plaque assay in MDCK cells at 36 hpi. Antibody dilutions of 0, 0.5, 1, 2.5 and 5 nM were prepared in FBS-free DMEM-GlutaMAX^TM^-I medium and mixed with the avicel overlay medium. Infection, chemical fixation and immunofluorescence staining of NP were carried out according to the above-mentioned protocol. R_INF_ [µm] and the relative reduction of R_INF_ in presence of increasing MEDI8852 concentration, normalized to control wells containing IAV infected cells without antibody were calculated and three-parameter nonlinear curve fits were generated in Prism 10.1.2 (GraphPad). From these fits, the IC50 (50% reduction of R_INF_) was obtained.

### Live cell imaging of IAV infected Calu-3 cells

For live cell imaging of fluorescent IAV plaques, Calu-3 cells infected with WSN:PAmScarlet or WSN-M1_Udorn_:PAmScarlet were overlaid with 1% (w/v) agarose (Biozym) overlay medium and imaged with the 5x objective of the CellDiscoverer 7 microscope (Zeiss) under constant conditions of 37°C, 20% O_2_ and 5% CO_2_. Cell nuclei were stained by addition of Hoechst (Sigma-Aldrich) into the overlay (1:1000 dilution). Images were acquired every 3 h over a course of 66 h, starting at 52 hpi or 79 hpi.

### Tracking and motion analysis of IAV infected Calu-3 cells

To analyze cell migration within IAV plaques, we performed tracking and motion analysis of Calu-3 cells. First, the global sample drift in the time-lapse fluorescence microscopy images was computed using a method based on cross-correlation [63,64] for the Hoechst channel. The determined drift was then corrected in both the Hoechst and the mScarlet channel by shifting the images. Second, the fluorescently labeled cells in the microscopy images were tracked using a probabilistic particle tracking method [65] based on multi-sensor data fusion and Bayesian filtering which combines Kalman filtering and particle filtering. Separate sensor models and sequential multi-sensor data fusion are used to integrate multiple detection-based and prediction-based measurements while considering different uncertainties. The measurements are obtained using elliptical sampling [66], and information from both past and future time points is exploited using Bayesian smoothing. Cell detection was performed by the spot-enhancing filter (SEF) [67] consisting of a Laplace-of-Gaussian (LoG) filter followed by intensity thresholding.

To analyze the motion of uninfected cells proximal to the infection plaque, a binary mask was created based on the mScarlet channel and computed trajectories within the mask were selected. First, segmentation of the infected cells in the mScarlet channel was performed by adaptive thresholding using a threshold defined by the mean intensity of the filtered image plus a factor times the standard deviation. The same factor was used for all images of an image sequence. Then, a binary mask of the infection plaque was created from the segmentation result using 40 iterations of binary dilation followed by hole filling using binary dilations and computing the binary sum over the complete image sequence. The largest connected component of the resulting binary mask was further dilated 120 times to create a mask for infection proximity. The final binary mask for analyzing infection-proximal uninfected cells was obtained by binary image subtraction of the infection mask from the infection-proximal mask. Computed trajectories in the Hoechst channel with most positions inside this mask were used for further analysis (Suppl. Figure 5). Additionally, the center of infection of an image sequence was determined as the center-of-mass of the segmentation result of the last image in the mScarlet channel. For uninfected control cells, the image center was used instead.

The motion of infected and uninfected cells was quantified by computing different motion properties from the computed trajectories. To improve the accuracy, only trajectories with a minimum duration of 12 hours (corresponding to 4 time steps) were considered. The velocity over time for each trajectory was calculated and then averaged for infection-proximal uninfected cells as well as for cells in the mScarlet channel and cells in the Hoechst channel of image sequences without infection plaques. The distance migrated of a trajectory was computed as the distance between the first and last position, and the distance migrated towards the center was calculated as the difference between the distance of the first position to the center of infection and the distance of the last position to the center. In addition, to characterize the movement of cells relative to the center of infection, we computed the cosine deviation as the cosine of the angle between the direction from the first position to the last position and the direction from the first position to the center of infection (Suppl. Figures 4 and 5). The values of the cosine deviation lie in the range -1 to 1 with positive values close to 1 indicating motion towards the center of infection, and negative values indicating motion away from the center. Since the cell density of infection plaques strongly increases over time, only the first 54 hours (corresponding to 19 time steps) were used to compute the above-mentioned motion properties to improve the accuracy. The motion properties were computed for each trajectory of an image sequence and then averaged over the image sequence.

### Entry time-course assay for spherical and filamentous IAV

To determine the entry half-time of spherical and filamentous IAV, A549 cells were seeded into 24-well plates (Corning) at a seeding density of 5×10^4^ cells/well and incubated for 24 h at 37°C and 5% CO_2_. Cells were infected on ice with spherical or filamentous fluorescent reporter virus at an MOI of 3 in FBS-free DMEM-F12-GlutaMAX^TM^-I to guarantee synchronized infection. After 1 h on ice, cells were washed three times with cold PBS. PBS was replaced by FBS-free DMEM-F12-GlutaMAX^TM^-I and cells were incubated at 37°C and 5% CO_2_. At 0, 5, 10, 20, and 60 minutes post infection (mpi), cells were treated with 50 mM NH_4_Cl to block endosomal acidification and subsequently incubated at 37°C and 5% CO_2._ At 12 hpi, cells were fixed with 4% PFA in PBS for 30 min at RT. Immunolabelling of IAV-NP and DAPI staining was performed according to the above-described protocol.

### IAV entry assay in presence of inhibitors

A549 cells were seeded into 24-well plates at a seeding density of 8×10^4^ cells/well and incubated for 24 h at 37°C and 5% CO_2_. Cells were incubated with different drug concentrations of Dynasore (324410, Sigma-Aldrich) (10, 50, 100 µM) or EIPA (A3085, Sigma-Aldrich) (40, 60, 80 µM) or DMSO in infection medium consisting of FBS-free DMEM-F12-GlutaMAX^TM^-I supplemented with 50 mM HEPES, 0.2 % BSA for 1h at 37°C and 5% CO_2_. Synchronized infection with spherical or filamentous WSN was performed on ice for 1 h at MOIs of 3 or 10, diluted in infection medium with inhibitor or DMSO. After virus attachment, virus inoculum was removed, and cells were washed 2 times with the corresponding cold inhibitor mix in infection medium. Plates were incubated with inhibitors at 37° and 5% CO_2_ for 6h, followed by washing with PBS and fixation with 4% PFA for 15 min at RT. Immunolabelling of IAV-NP and DAPI staining was performed according to the above-described protocol. Fluorescence microscopy images were acquired using the 20x objective of the CellDiscoverer 7 microscope (Zeiss) (pixel size 0.3450×0.3450 µm^2^, 16 bit depth). Quantification of IAV infection for entry assays was performed in ImageJ/FIJI [58], using the StarDist plugin [60] to define regions of interest (ROIs). An intensity threshold of NP signal within these ROIs was set and cells with an average intensity above the threshold were counted as infected. Experiments were performed in three independent replicates.

### Sample preparation for cryo-electron microscopy

Glow-discharged gold EM grids (200 mesh, Au, holey SiO, R1/2, Quantifoil) were placed into polydimethylsiloxane Sylgard 184 (Farnell GmbH) coated 35 mm dish (Corning) [68] and disinfected with 70% ethanol for 30 min. The dish was washed three times with DMEM-F12-GlutaMAX^TM^-I 10% FBS and 1% P/S. A549 cells were seeded at a density of 1.8 × 10^5^ cells per dish and incubated at 37°C and 5% CO_2_ for 24 h. Virus was thawed on ice for 1 h and a dilution of 5 × 10^6^ PFU/ml was prepared in FBS-free DMEM-F12-GlutaMAX^TM^-I. 20 µl of virus dilution were pipetted onto parafilm placed in a 10 cm dish on a cooling plate at 4°C. EM-grids were blotted on Whatman No.1 filter paper, placed onto the drop of virus and incubated for 30 min. Grids were washed 3 times with FBS-free DMEM-F12-GlutaMAX^TM^-I and placed in the incubator at 37°C for 15-30 min. Cells were immediately plunge-frozen into liquid ethane using the following settings of an automatic EM GP2 plunge-freezing device (Leica): chamber temperature: 37°C, humidity: 80%, back-side blotting: 3.5 sec. Grids were stored in LN_2_ until further processing.

### Cryo-focused ion beam milling (cryo-FIB milling)

Cryo-lamellae of infected cells were prepared by cryo-focused ion beam milling with an Aquilos2 dual-beam cryo-focused ion beam-scanning electron microscope (cryo-FIB-SEM, ThermoFisher Scientific) with a cryo-stage at 180°C. Grids were sputter coated before and after deposition of an organometallic platinum layer via the Gas Injection System (GIS). Using a gallium ion beam at an acceleration voltage of 30 keV, cells were semi-automatically milled at a stage angle of 15°. Three steps were milled using an adapted protocol for automated cryo-lamella preparation, including micro-expansion joints [69,70]. The last two polishing-steps were performed manually to achieve a nominal lamella thickness of 150 nm.

### Cryo-electron tomography of IAV infected cells

Cryo-ET data of lamellae from IAV infected A549 cells were acquired with SerialEM [53] using a Titan Krios cryo-TEM (ThermoFisher Scientific) operated at 300 keV and equipped with a Quanta Imaging Filter (Gatan) with an energy filter slit set to 20 eV and a K3 direct electron detector (Gatan). Grids were first mapped at 8,700 x magnification (pixel spacing: 10.64 Å). Endosomes containing viruses were selected for tilt series acquisition at 33,000 x magnification (pixel spacing: 2.67 Å) using PACEtomo [55] in SerialEM following a dose-symmetric tilting scheme [56] and the following settings: zero angle set to 8°, nominal tilt range from +68° to -52° and 3° increments, target defocus -3 µm, electron dose per record 3e^-^/Å^2^. Drift correction of acquired movies was done with Motioncor2 [57]. Tomographic reconstruction was performed with AreTomo [71]. 3D segmentation was done using the Dragonfly software (Version 2022.2.0.12227, Comet Technologies Canada).

## Acknowledgements

We thank Dr. Tijana Ivanovic for kindly providing MEDI8852 antibody and Dr. Michael D. Vahey for kindly providing a plasmid pCDNA3.1RG-WSN-M1_Udorn_M2_WSN_. We thank the Infectious Diseases Imaging Platform (IDIP) at the Center for Integrative Infectious Disease Research Heidelberg and the cryo-EM network at Heidelberg University (HD-cryoNET) for support and assistance. The authors would like to thank the Soft (bio)materials characterisation Core Facility (Biomechanics) at IMSEAM for their support. The authors gratefully acknowledge the data storage service SDS@hd supported by the Ministry of Science, Research, and the Arts Baden-Württemberg (MWK), the German Research Foundation (DFG) through grant INST 35/1314-1 FUGG and INST 35/1503-1 FUGG.

## Declarations

### Author contributions

Conceptualization: SP, CL, CS, JM and PC; Methodology: SP, CL, CS, JM, XP, VL, MV, LE, SK, FG, KR; Investigation: SP, CL, CS, JM, XP, VL, MV, LE; Visualization: CL, CS, JM; Funding acquisition: CF, EAC-A, CS-U, KR, and PC; Project administration: PC; Supervision: CF, EAC-A, CS-U, KR, and PC; Writing - original draft: SP, PC; Writing - review & editing: all

### Funding

This work was supported by a research grant from the Chica and Heinz Schaller Foundation (Schaller Research Group Leader Programme) and by the Deutsche Forschungsgemeinschaft (DFG, German Research Foundation): CF, EAC-A, CS-U, KR, and PC project no. 240245660-SFB1129; PC: 437060729. M.V. and CS-U were supported by the Max Planck School “Matter to Life” funded by the German Federal Ministry of Education and Research (BMBF) and the Flagship Initiative “Engineering Molecular Systems”. We also acknowledge funding from the DFG under the German Excellence Strategy 2082/1-390761711 (3D Matter Made to Order) and the European Research Council (ERC CoG no. 101001797 PHOTOMECH).

### Disclosure and competing interest statement

The authors declare that they have no conflict of interest.

### Data availability

Electron tomography data will be deposited to EMDB and will be available upon publication. Additional data and material related to this publication may be obtained upon request.

## Notes

### Competing Interest Statement

The authors have declared no competing interest.

