## Supplementary data for "Morphology-dependent entry kinetics and spread of influenza A virus"

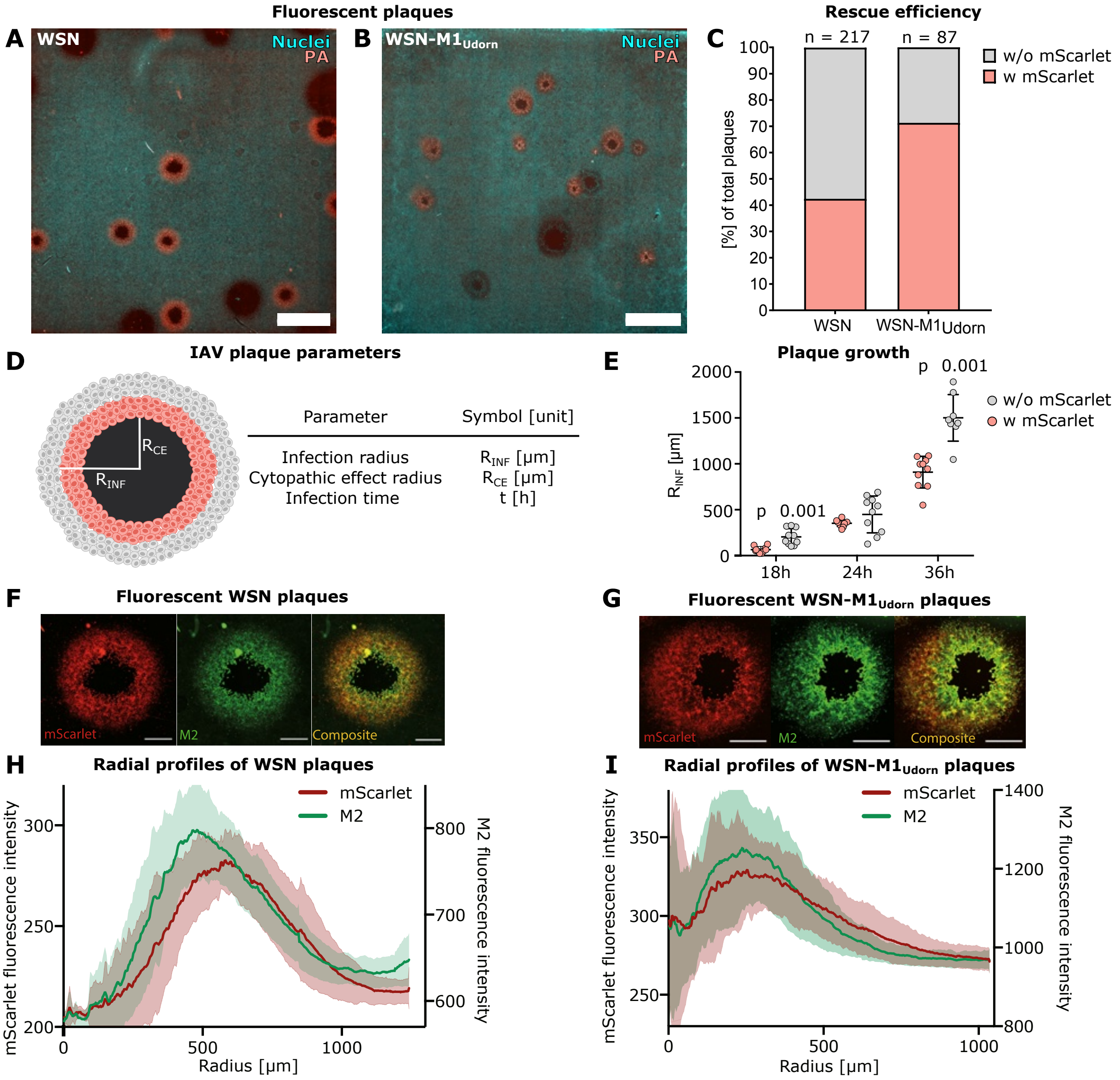


**Supplementary Figure 1: Characterization of fluorescent IAV plaques. (A)** Exemplary images of plaques from MDCK cells infected with WSN:PAmScarlet or **(B)** WSN-M1$_{Udorn}$:PAmScarlet at 36 hpi with PA:mScarlet (red) and nucleus staining (cyan). Scale bars 3 mm. **(C)** Percentages of plaques with (w) (red) and without (w/o) (grey) mScarlet signal for WSN:PAmScarlet (n=217) and WSN-M1$_{Udorn}$:PAmScarlet (n=87), quantified at different timepoints. **(D)** Schematic representation of the zones within a fluorescent plaque and parameters for quantification of IAV cell-to-cell spread. **(E)** Infection radius ($R_{INF}$) of plaques with (red) and without (grey) mScarlet quantified for 10 plaques per condition, at 18, 24, 36 hpi. Exemplary images of a fluorescent plaques in MDCK cells with PAmScarlet (red) and immunostained M2 (M2) and composite of both signals for **(F)** WSN:PAmScarlet and **(G)** WSN-M1$_{Udorn}$:PAmScarlet. Scale bars: 500 µm. Mean fluorescence intensities of PAmScarlet (red) and M2 (green) from radial profiles of 10 plaques for **(H)** WSN:PAmScarlet and **(I)** WSN-M1$_{Udorn}$:PAmScarlet. Standard deviations are indicated.

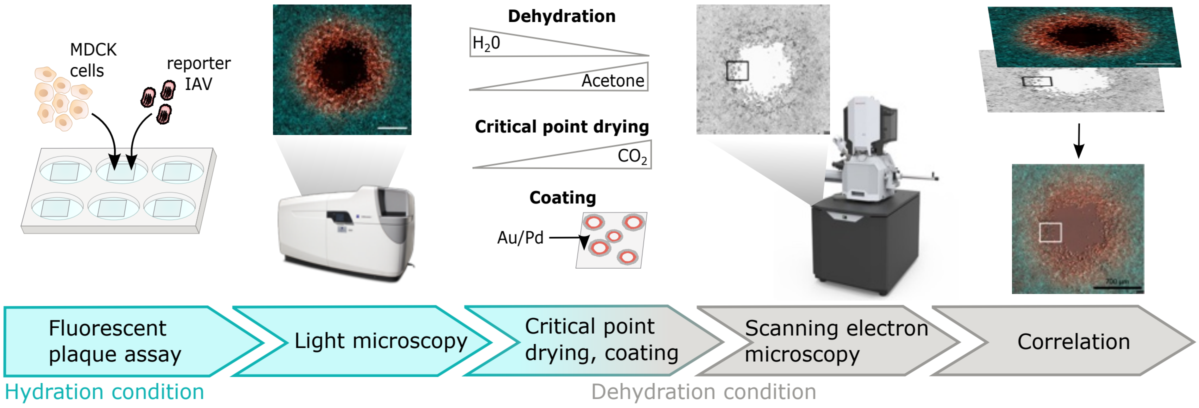


**Supplementary Figure 2: Workflow of correlative light and scanning electron microscopy for the study of IAV plaque growth.** MDCK cells were seeded onto ITO-coated coverslips and infected with spherical or filamentous reporter IAV expressing PAmScarlet. Plaques were imaged by fluorescence microscopy. Samples were prepared for scanning electron microscopy (SEM) by dehydration with increasing acetone concentrations. Critical point drying and sputter coating with Au/Pd was performed. SEM overview maps of plaques were acquired for correlation with fluorescent images. High magnification SEM images were acquired.

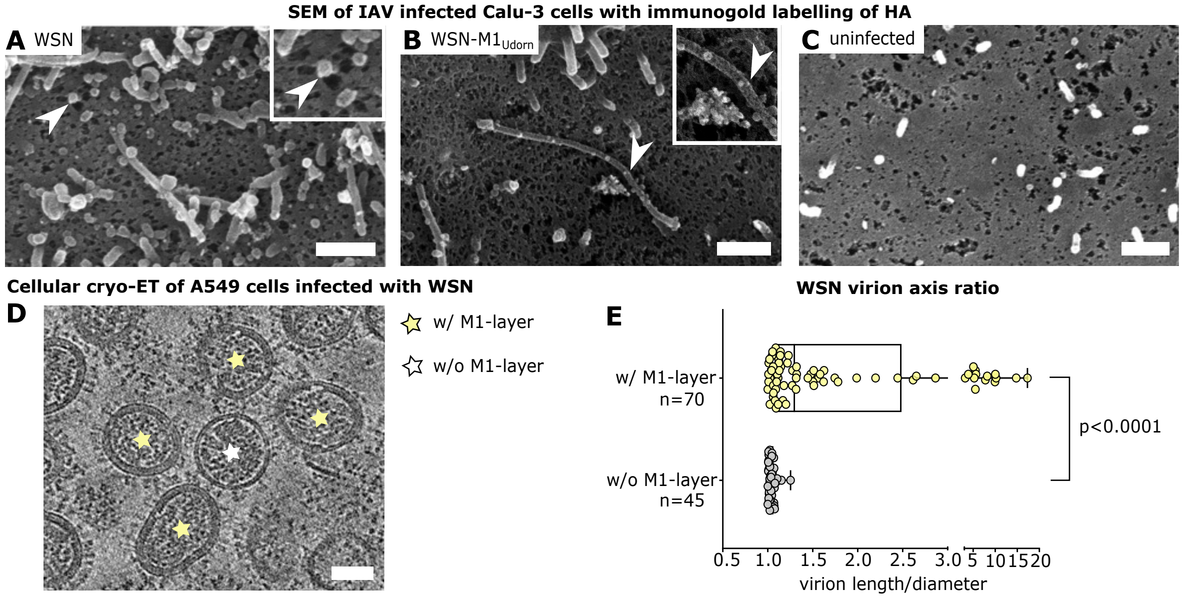


**Supplementary Figure 3: Immunoelectron microscopy of IAV plaques and virion morphology during cell entry.** Scanning electron microscopy (SEM) of Calu-3 cells infected with **(A)** WSN:PAmScarlet, **(B)** WSN-$M1_{Udorn}$:PAmScarlet, fixed at 4 days post infection. White arrowheads indicate 20 nm immunogold labelling of hemagglutinin (HA). **(C)** SEM of uninfected Calu-3 cells. Scale bars 500 nm. **(D)** Slices through a cryo-electron tomogram showing spherical WSN particles in endosomal compartments of infected A549 cells at 15-30 min post infection. Virions with assembled matrix protein 1 layer (w/ M1-layer) are highlighted with yellow stars. One virion without (w/o) M1-layer is highlighted with a white yellow star. **(E)** WSN virion length/diameter ratio of particles w/ M1-layer (yellow) and w/o M1-layer (grey) within endosomes of infected A549 cells, quantified from cryo-electron tomograms. Significance analysis was done by unpaired t-test.

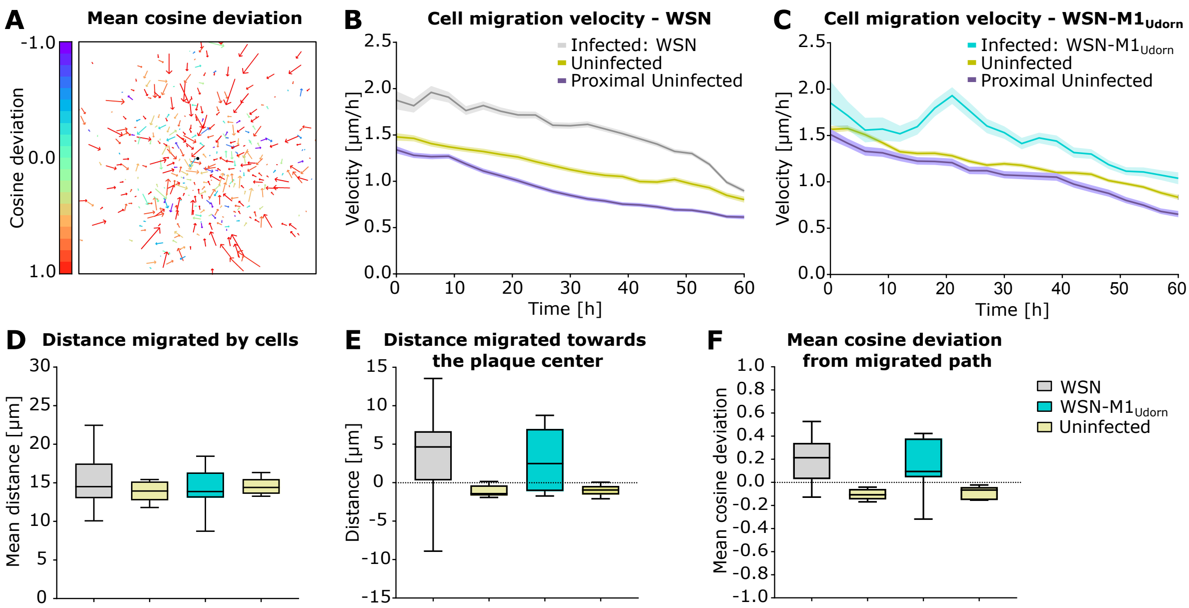


**Supplementary Figure 4: Distance, velocity and direction of Calu-3 cell migration. (A)** Migrated paths of Calu-3 cells tracked from their first to last position of time-lapse movies. Color code of arrows represents the cosine deviation from migrated path, where +1 indicates migration to the plaque center and -1 indicates migration away from the center. **(B, C)** Mean velocity of cell migration trajectories for infected and uninfected Calu-3 cells. Standard errors are indicated. The time point 0 h corresponds to 52 h post infection. **(D)** Mean migrated distance of Calu-3 cells from their first to last position. Boxes show quartiles with median lines and min to max values. **(E)** Migrated distance of Calu-3 cells towards the plaque center. For each movie, the distances are averaged over all trajectories. **(F)** Mean cosine deviation of cell movement from migrated path with positive values indicating migration towards the center of infection. Color code of plots B-F: grey: infected with WSN:PAmScarlet (spherical), cyan: infected with WSN-M1$_{Udorn}$:PAmScarlet (filamentous), yellow: uninfected cells, blue: proximal uninfected cells.

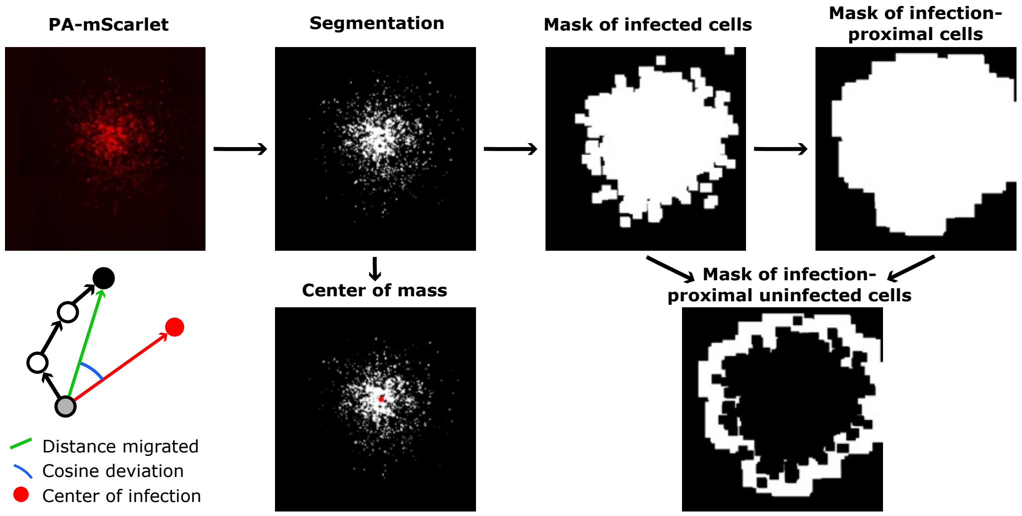


**Supplementary Fig. 5: Workflow of tracking and motion analysis for cells within IAV foci.** IAV infected cells expressing PA-mScarlet (red) were segmented by adaptive thresholding. From the last segmented image of the time-lapse series, the center of infection was determined as the center of mass. Based on the segmentation, binary masks of infection foci were created by binary dilation and hole filling. A mask for infection-proximal cells was created by further dilating this mask 120 times. To obtain a mask for uninfected cells in proximity to IAV foci the mask of infected cells was subtracted from the mask of infection-proximal cells. Cell migration was determined by probabilistic particles tracking. The distance migrated from the first to the last position (green) and the cosine of the angle (blue) between the directions from the first to the last position (green) and the first position and the plaque center (red) were computed.
